# ALANizer: Design and validation of experimental lighting rig for studying artificial light at night in ecosystems

**DOI:** 10.1101/2025.08.30.673291

**Authors:** Daphne Chevalier, Nisa Chavez Molina, Juli Carrillo, Quentin Geissmann

## Abstract

Over the past few centuries, human activity has introduced astoundingly bright light to the environment at a rapidly accelerating pace. Artificial light at night (ALAN) has detrimental effects on a wide range of organisms, including arthropods. To minimise impact, we must understand how different organisms are affected and whether certain lights are less perturbing. We designed a novel tool for introducing and monitoring different types of ALAN, called ALANizer, composed of open-source software and inexpensive, widely-available hardware. We describe the design and assembly of ALANizer as well as a feasibility study where we installed 12 ALANizers in hedgerows and monitored arthropod activity using pitfall traps. Understanding ALAN’s impacts on ecosystems only grows more pressing as LED technology proliferates. As an effective, low-cost, and scalable tool for introducing and monitoring ALAN in ecosystems, ALANizer can help advance this key area of research.

## 2 Introduction

To protect biodiversity, we urgently need to better understand the effects of artificial light at night (ALAN) on ecosystems (Hölker et al., 2021). Despite copious evidence linking ALAN to negative biological impacts on individual species, little is known about its role in disrupting ecological communities; meanwhile, ALAN continues to intensify and spread (Hölker et al., 2021). Emerging recommendations suggest that amber-coloured LEDs may be less harmful than white-coloured LEDs, but supporting evidence is lacking, especially for arthropods (Jägerbrand & Bouroussis, 2021).

Experimental ecology requires controlled perturbations of the natural environment that can be monitored with effects quantified (Clobert et al., 2018). Although the scientific community has developed effective methods for manipulating many environmental variables, there is not yet a standardised and widely-available approach for introducing artificial light to field settings (Jägerbrand & Bouroussis, 2021). Studies featuring existing light installations, such as streetlights, can be useful, yet are limited by confounding variables, such as noise, heat, and increased human activity, and primarily provide observational data, which precludes causal links (Owens & Lewis, 2018). Correlative approaches also require equipment to measure important covariates, such as light intensity.

It is difficult to know to what extent ALAN has already affected ecosystems and biodiversity. However, we can perform field experiments in areas that are still relatively ALAN-naïve (without histories of artificial lighting), introducing light at a small scale and observing ecological consequences. Although some groups have designed and deployed mobile experimental lighting rigs in these settings, costly setups limit the number of experimental replicates (Boyes, 2021), and suitable experimental sites often lack the required infrastructure. For example, the setup described by Lang et al. (2025) is designed for connection to a power grid. Design for battery-powered operation is an important consideration for studying ALAN in historically dark areas.

Experimental lighting rigs must introduce light on a schedule, vary their colour and/or intensity, and monitor lighting conditions in the environment. Most light metrics, especially those derived from lumens, are tailored to human vision. In ecological research, it is especially important to measure light from the perspectives of other vision systems (Aulsebrook et al., 2022). Consequently, some researchers have suggested concurrent monitoring of intensity and spectral characteristics (Seymoure et al., 2023; Grubisic et al., 2018).

In this article, we address the need for an autonomous, scalable, and controllable device to study ALAN in the field. First, we describe ALANizer, a novel open-hardware and open-source tool. Then, we present a sample deployment of ALANizer to illustrate its usefulness for exploring ALAN’s effects on terrestrial arthropods.

## 3 Application overview

ALANizer consists of hardware and firmware components, whose operation and performance were validated in-lab and experimentally. First, we detail ALANizer’s hardware components (Fig. 1). We then describe firmware components, including programs to schedule the light timing and to collect data from the light sensors. We report validation steps and results (Fig. 2). Finally, we describe an experimental deployment in real-world field conditions (Fig. 3).

**Figure 1:**
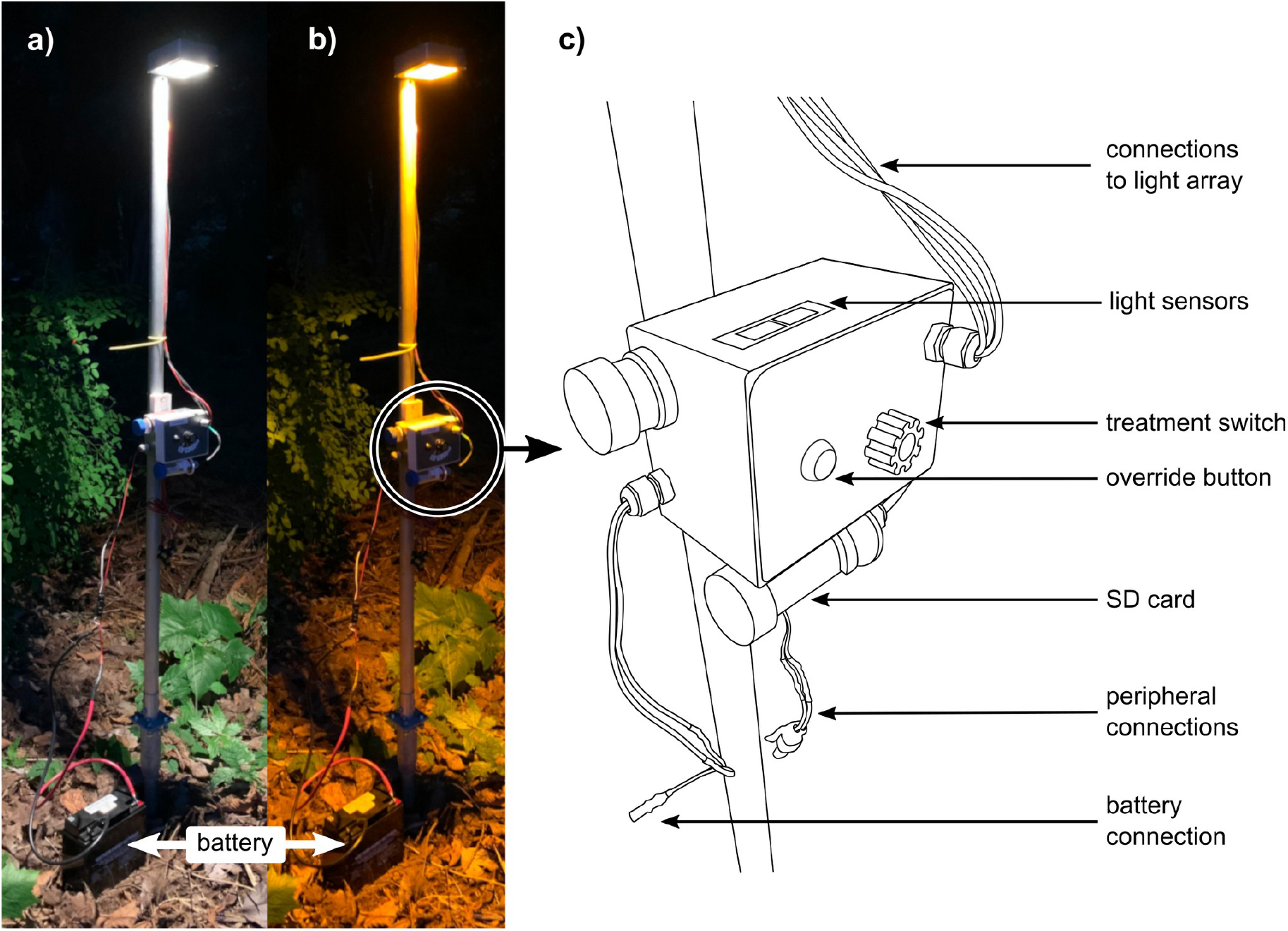
ALANizer overview. Deployed device can be set to white-light treatment (a) and amber-light treatment (b), displayed alongside diagram of ALANizer’s controller box (c). The treatment switch toggles between unlit, white-light, and amber-light treatments. The override button powers the light array (if the treatment switch is set to a lit treatment) regardless of schedule, which allows the user to check that the device is outputting the correct treatment at any time. The SD card saves light sensor data every 5 minutes. It is stored in an external tube and can be removed without opening the controller box. The peripheral connections can power other sensors or tools.

**Figure 2:**
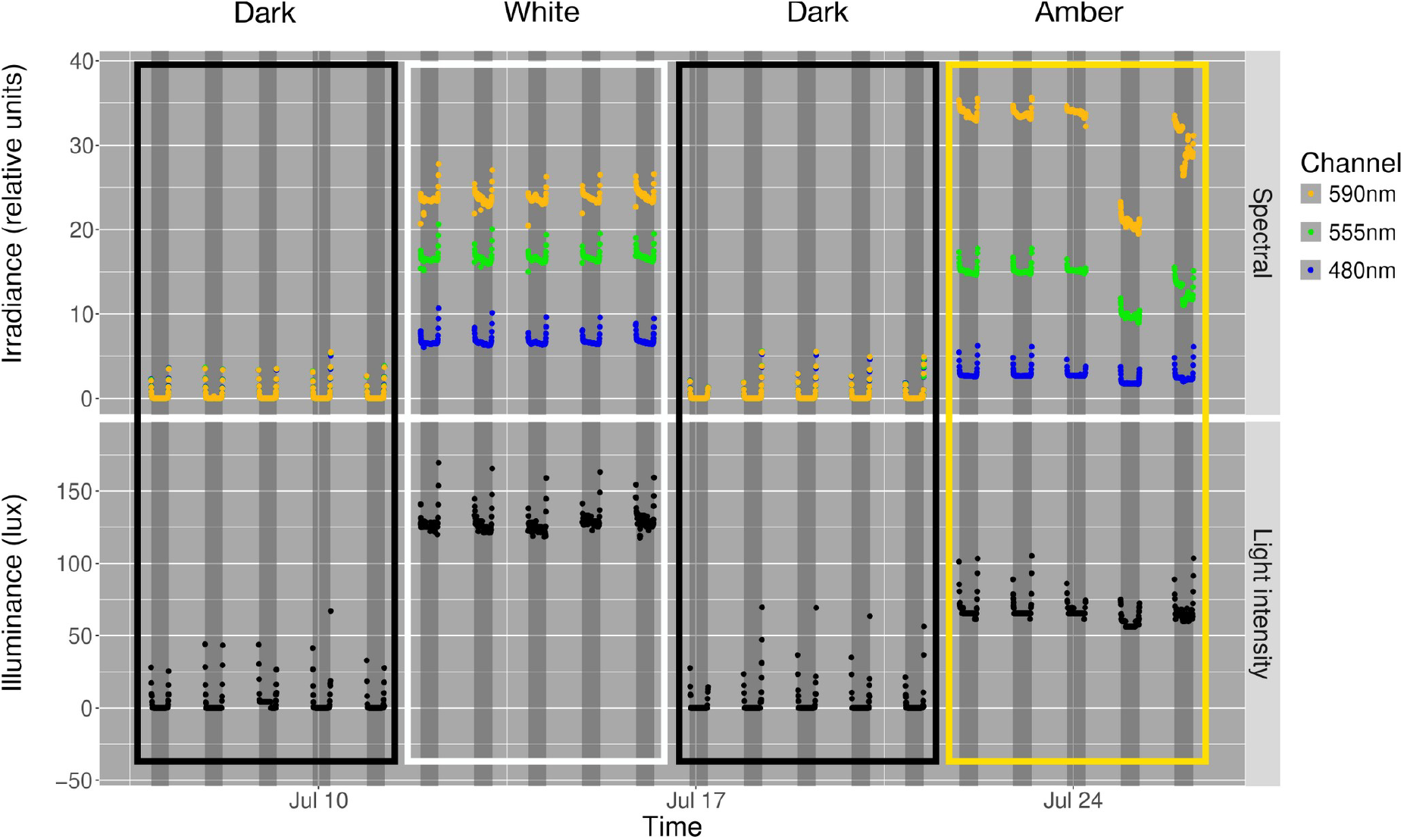
Example light sensor measurements from ALANizer. Plots of nighttime spectral and intensity measurements as light is turned on and off in a scheduled manner, for white/amber/control settings (taken from Site 6). Irradiance units are relative while illuminance is displayed in lux. Dark grey bands represent civil night (i.e., when the sun is ≥ 6 degrees below the horizon). We measured very little light during dark settings. The white treatment exhibited more green (555nm) and blue (480nm) light and higher overall light intensity than amber light.

In this section, we detail ALANizer’s specifications, components, operation, and validation. We identified several key specifications for our system by reviewing a range of ecological studies centred around ALAN. Justifications for these benchmarks can be found in “Components,” below.

We wanted ALANizer to operate off-grid (through battery power) with enough capacity to produce light over at least five consecutive nights, and emit light in two colours (white and amber), at a luminous flux of approximately 1000 lumens (equivalent to a 75 watt incandescent bulb). Additional needs included the ability to schedule when ALANizer’s light turned on and off each day, in advance of deployment, and the ability to monitor overall light intensity and spectral character between 350 and 1000 nanometres. These data needed to be retrievable without exposing the device’s electronics to the elements. Lastly, ALANizer needed to be robust and weatherproof, keeping internal components dry during months outside.

## 4 Components

### 4.1 Hardware overview

ALANizer (illustrated in Fig. 1) hardware allows users to introduce and monitor light in the field. This is done by supplying power to the unit (“power source”, see “Supplementary materials”) which activates the experimental light source (“light box”) which can be cycled through light settings and measured via the “controller box.” We provide detailed schematics on the ALANizer GitHub repository (see “Supplementary materials”).

#### 4.1.1 Light box

The ALANizer light box must emulate realistic light sources for both amber and white lights, be low-power enough to rely on batteries, and operate without producing unmanageable heat. We therefore designed a printed circuit board (PCB) with two parallel strings of high-efficiency LEDs (Cree XP-E2) in series, connected to a low-side treatment switch (Cree LED, 2024). Although LEDs are more efficient than other lamp types, high-intensity LEDs require thermal management. The LED array PCB incorporates many thermal vias and a solid copper pour on the back. In our implementation, we bolted the PCB to an aluminum enclosure with a layer of thermal paste between the solid copper pour and enclosure, turning the enclosure into a large heat sink.

We measured illuminance at ground level (with lights 150cm from ground) with a Lambda Photometric 2219-7903 sensor combined with a LI-COR LI-185A reader (LI-COR, n.d.; Lambda Instrument Co., n.d.). Intensity directly underneath the lights was recorded as approximately 44 lux and 43 lux at ground level under the white and amber settings respectively, and 1 lux for both light settings at a distance of 5 metres away.

#### 4.1.2 Controller box

The controller box contains hardware for scheduling, adjusting, and monitoring the light treatment. We used an Arduino Mega as the controller, which interfaced with two light sensors (and, optionally, other peripherals). We connected an external real-time clock (RTC) circuit to keep time, which allowed us to schedule the light’s on and off periods and to timestamp sensor data.

We wanted to monitor both light intensity and light colour (spectral character) following best practices suggested by Seymoure et al. (2023). We integrated a light intensity sensor (Adafruit TSL2591, which measures wavelengths between 400 and 1100 nm, with a dynamic range of 188 µlux-88,000 lux) and a spectral sensor (Adafruit AS7341, which measures wavelengths between 415 and 680 nm) into the enclosure, pointing upward toward the light. We mounted a piece of frosted acrylic over the sensors as a diffuser, reducing error by scattering incoming light in a uniform manner (UDT Instruments, 2020).

We added a rotary switch to allow the user to toggle between the light’s different states, which are “control” (light off), white light treatment, and amber light treatment. We also added an override switch, through which the user can override schedule and turn the light on during the daytime, which is useful for ensuring the device works.

To keep the SD card accessible, we placed the SD module in a 50mL centrifuge tube (FalconTM) attached to the outside of the controller box (see “Supporting information”). We replaced the tube’s conical tip with a 3D-printed piece, securely attached with “gasket maker” (Permatex). This piece’s integrated cable gland kept the SD module connected to the controller while maintaining a watertight seal. The tube’s threaded lid provided access to the SD card.

### 4.2 Firmware

ALANizer’s firmware can be separated into pre-deployment and post-deployment utilities. Before deployment, the user needs to generate a list of timestamps for turning the light on and off based on sunset and sunrise times given geolocation and time of year, which is then stored on the SD card (see “Initialisation and setup” for instructions). The user also needs to synchronise all RTCs to real-time (see the Github repository for details).

After deployment, ALANizer’s controller collects sensor data every 5 minutes and stores it on an SD card. The controller also turns on an LED in the enclosure if it enters an error state (e.g., cannot communicate with a peripheral). The controller saves sensor data into a new file every hour, which prevents significant data loss from interrupted transmission but requires the user to later concatenate all data.

## 5 Operation

As mentioned in the firmware section above, operating ALANizer involves a few initialisation steps, followed by regular maintenance.

The ALANizer’s Arduino microcontroller must be programmed with the length of each day and night between light-switching events, which will be specific to season and location. We used the `suncalc` package in R (see “`scheduling.R`” in the ALANizer Github repository) (Thieurmel & Elmarhraoui, 2022; R Core Team, 2021).

To retrieve data, we replaced the SD cards each cycle and backed them up using `robocopy` or `rsync`. During these visits, we also replaced system batteries.

## 6 Validation

We validated ALANizer via several in-lab unit tests and a longer-term field deployment, which are detailed below.

### 6.1 Quality control (in-lab)

We tested the devices’ waterproofness by standardised engineering procedure (see “Supplementary materials”). We then tested basic functionality to ensure the sensors, data collection and storage, light scheduling, and toggling between settings functioned as expected.

### 6.2 Experimental deployment

To validate ALANizer’s functionalities, we introduced ALANizer into the environment and quantified differences in terrestrial arthropod activity dependent on experimental light conditions, across 12 sites. We chose sites located along hedgerows at an experimental research farm at the University of British Columbia’s Vancouver campus, at least 50 metres apart. We deployed an ALANizer at each site from the end of June to the end of November, 2023. Results from this study are available in Chevalier et al., 2025 (in prep).

During deployment, each ALANizer cycled through treatment settings (white light, amber light, or no light (control)) in a 20-day sequence split into four phases, where each phase lasted 5 days. All the lights followed the same pattern, where lit phases alternated with control phases, switching between white and amber treatments during alternating lit phases. We staggered schedules for different sites, so that at any given time, six sites were set to control, three were set to the white treatment, and three were set to the amber treatment.

As expected, light intensity was higher in lit phases than in dark control phases and light colour changed depending on whether the white or amber light was on. We observed the fluctuations shown in Figure 2, which matched our expectations, confirming that ALANizer introduced distinct wavelengths to the environment according to our experimental cycle.

## 7 Discussion

We built ALANizer, a novel and open-source tool that contributes to the ALAN research community by allowing researchers to introduce inexpensive and scheduled light to an experiment. Its core components cost less than $200 CAD and are highly adaptable to a range of deployment contexts (light designs, sensor needs, sunrise and sunset times, etc.).

Using ALANizer, we successfully introduced white and amber light to a light-naïve environment while collecting data on light levels and arthropod activity. ALANizer successfully recorded higher levels of blue and green light under the white treatment condition relative to the amber treatment (Fig. 2). Additionally, we found that the light intensity of our white treatment condition matched that of a standard streetlight (City of North Vancouver, 2024).

We identified several areas of improvement for ALANizer. Currently, ALANizer lacks wireless data transmission, requiring regular visits to experimental sites to download data. This may not be an issue for off-grid deployments, where batteries regularly need to be changed. However, it would be helpful for notifying the user when sensors are unexpectedly obstructed (i.e. by a fallen leaf), or when a peripheral device is unexpectedly disconnected. Additionally, retrieving and replacing the SD card without introducing water to the SD module can be challenging in heavy rain.

Many researchers have proposed guidelines and ideas for future ALAN studies (Jägerbrand & Bouroussis, 2021). These studies may be optimised by using lighting setups that are representative of real systems designed for human use. Future ALAN field experiments would benefit from assessing whether the lights being studied provide sufficient illumination for humans while mitigating effects on other organisms.

ALANizer’s low price, open-source documentation, and scalability make it well-suited to experimental ecology, which often requires many replicates to overcome the potential confounding effects of uncontrollable environmental variables.

ALAN’s acceleration shows no sign of stopping, making mitigation efforts increasingly crucial. To build effective policies and guidelines, we need high-quality evidence about its impacts, which we know to be wide-ranging and profound. This work establishes ALANizer as a promising new tool in the rapidly growing field of ALAN research, facilitating further investigation of ecological impacts from new lighting installations.

## 8 Statement on inclusion

Our study was performed on the traditional and ancestral territory of the x^w^məθk^w^əý əm (Musqueam) people. Our study involved input and participation from scientists and farm workers who work and live on and around the study area.

## 9 Author contributions

Daphne Chevalier, Juli Carrillo, and Quentin Geissmann conceived the ideas and designed methodology, with insight from Nisa Chavez Molina on pitfall trap placement. Daphne Chevalier and Nisa Chavez Molina collected the data; Daphne Chevalier, Juli Carrillo, and Quentin Geissmann analysed the data; Daphne Chevalier led the writing of the manuscript. All authors contributed critically to the drafts and gave final approval for publication.

