## Supplementary Materials for "ALANizer: Design and validation of experimental lighting rig for studying artificial light at night in ecosystems"

### 1 Resources

Documentation: https://github.com/chevalierid/alanizer/

Anonymised data: https://zenodo.org/records/15170657

### 2 Power supply

The ALANizer's power source must supply power to all of its components, over a sufficiently long experimental period to be biologically relevant and logistically feasible. The ALANizer's light consumes the majority of the device's overall power requirement, so the desired brightness and length of a given deployment period are determining factors in power supply selection. It is extremely important to respect depth-of-discharge limits: even deep-cycle marine batteries, which can be depleted to a greater extent than others, should be discharged by no more than 80% [1]. Using a maximum night length of 16 hours and our light array's current draw of one ampere, we chose a 300 amp-hour battery to run ALANizer for five nights, as seen in the calculations below.

5 LEDs in series:

52.8V * 1A = 15.6W

The Pololu U3V70A converts 6V battery to 15V with approximately 90% efficiency [2].

(battery voltage) * (current drawn from battery) * 0.9 = (sum of voltage across lights) (current across all lights)

$V_{bat} * I_{in} * 0.9 = 15.6V * 1A$

$6V * I_{in} * 0.9=15.6V * 1A$

$I_{in} = 15.6 / (0.96)$

$I_{in} = 2.8A$

16h = max. night length

16h * 5 nights = 80h

2.8 * 80 = 224h

224 / 300 = 74.6% discharged

The power draw of high-intensity LEDs dictates many aspects of ALANizer setup. We chose deep-cycle marine batteries for their high capacity and physical durability. We used small batteries during dark control cycles. Fully sealed batteries are preferable to flooded batteries, since acid can leak during transit over uneven ground.

Within the ALANizer controller box, several key hardware pieces regulate and supply electrical power from the battery. There are two separate voltage regulators, one for the Arduino (which then supplies the clock, SD card reader, and sensors) and one to supply a constant-current LED driver that provides the LED array with power. We designed 3D-printed brackets to keep components immobile within the controller box.

### 3 Controller box design and waterproofing

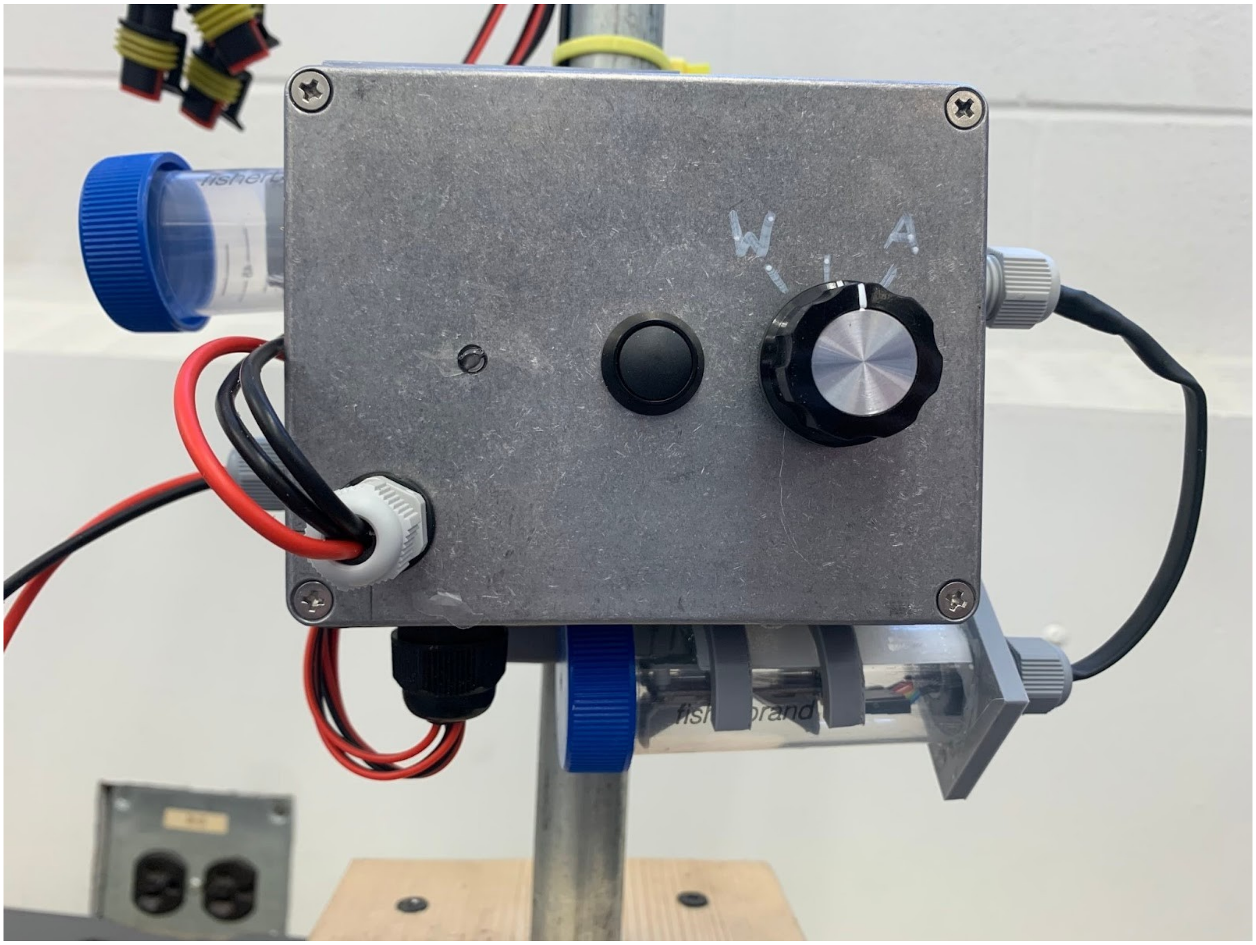

**Figure S.1: SD card location on ALANizer.** Modified centrifuge tube holding SD card breakout board visible at bottom right.

We took several precautions to maintain a watertight seal between the controller box’s contents and the exterior. We used NEMA-rated cable glands at all cable entry points. We also adhered the diffuser to the enclosure with gasket maker (Permatex). We applied epoxy resin to all 3D-printed parts.

Temperature and moisture inside both the light box and the controller box also fluctuated with environmental conditions. We placed desiccant bags (silica) within each enclosure to mitigate condensation. We found that a very large amount of water in an enclosure was more likely to be a sign of water ingress and inadequate waterproofing. To test waterproofness, we used a drip proof test described by the Radio Technical Commission for Aeronautics [3].

### 4 Site locations

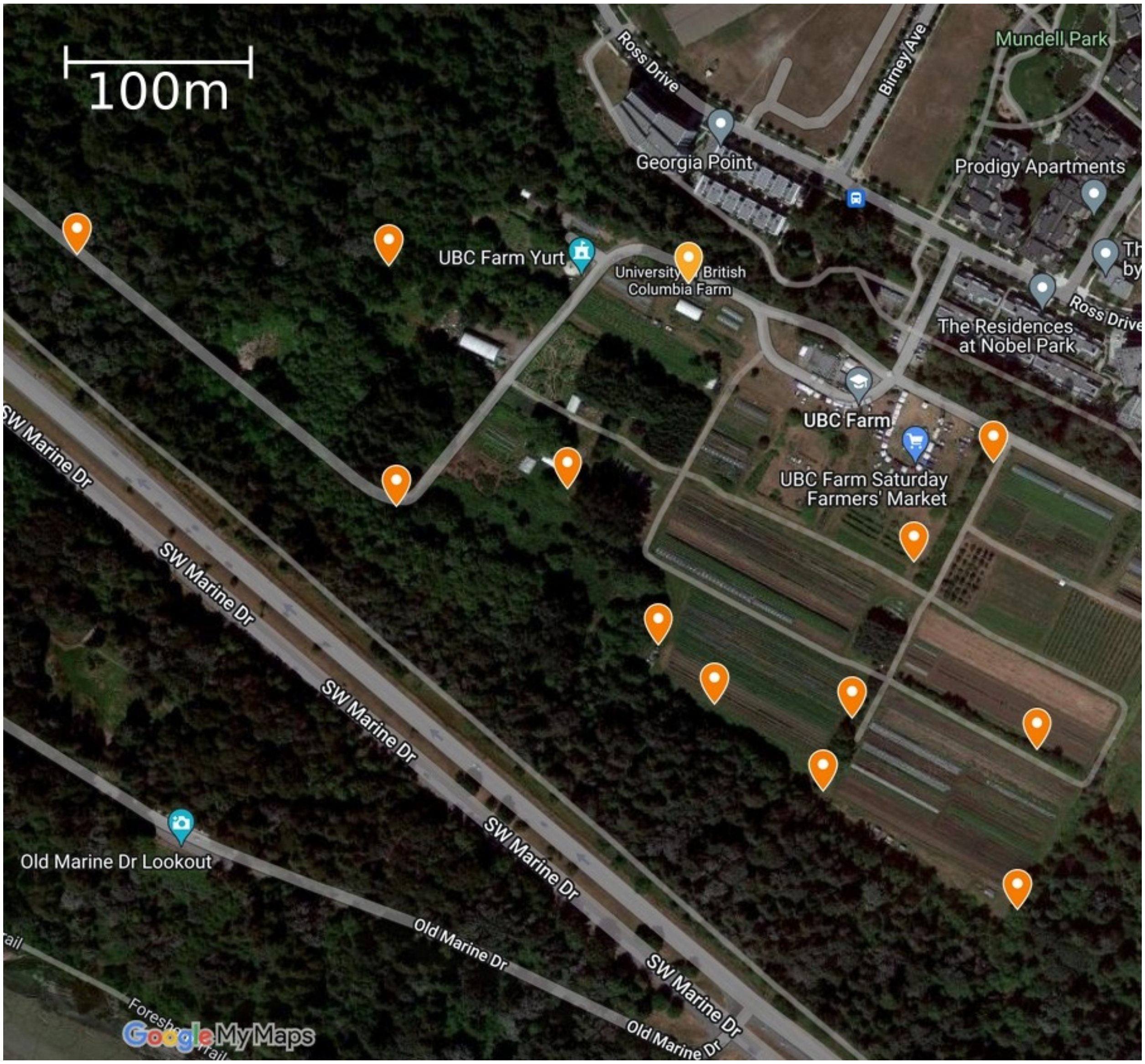


**Figure S.2: Site locations at the UBC Farm.**

### 5 References

[1] Century Yuasa Batteries Pty Ltd. (n.d.). Deep cycle batteries explained. *Battery Zone*. https://www.batteryzone.co.nz/the-zone/expert-advice/deep-cycle-batteries-explained

[2] Pololu Robotics & Electronics (n.d.). 4.5-20V Fine-Adjust Step-Up Voltage Regulator U3V70A. *Pololu*. https://www.pololu.com/product/2890

[3] Radio Technical Commission for Aeronautics (2010). Environmental Conditions and Test Procedures for Airborne Equipment. GlobalSpec. https://standards.globalspec.com/std/9894152/rtca-do-160
